# Acute-phase CD4+ T cell responses targeting invariant viral regions are associated with control of live-attenuated simian immunodeficiency virus

**DOI:** 10.1101/321000

**Authors:** Matthew S. Sutton, Amy Ellis-Connell, Ryan V. Moriarty, Alexis J. Balgeman, Dane Gellerup, Gabrielle Barry, Andrea M. Weiler, Thomas C. Friedrich, Shelby L. O’Connor

**Author notes:** Address correspondence to Shelby L. O’Connor.

## Abstract

We manipulated SIVmac239Δnef, a model of MHC-independent viral control, to evaluate characteristics of effective cellular responses mounted by Mauritian cynomolgus macaques (MCMs) who express the M3 MHC haplotype that has been associated with poor control of pathogenic SIV. We created SIVΔnef-8x to test the hypothesis that effective SIV-specific T cell responses targeting invariant viral regions can emerge in the absence of immunodominant CD8+ T cell responses targeting variable epitopes, and that control is achievable in individuals lacking known protective MHC alleles. Full proteome IFNγ ELISPOT assays identified six newly targeted immunogenic regions following SIVΔnef-8x infection of M3/M3 MCMs. We deep sequenced circulating virus and found that four of the six newly targeted regions rarely accumulated mutations. Six animals infected with SIVΔnef-8x targeted at least one of the four invariant regions and had a lower set point viral load compared to two animals that did not target any invariant regions. We found that MHC class II molecules restricted all four of the invariant peptide regions, while the two variable regions were restricted by MHC class I molecules. Therefore, in the absence of immunodominant CD8+ T cell responses that target variable regions during SIVmac239Δnef infection, individuals without ‘protective’ MHC alleles developed predominantly CD4+ T cell responses specific for invariant regions that may improve control of virus replication. Our results provide some evidence that antiviral CD4+ T cells during acute SIV infection can contribute to effective viral control and should be considered in strategies to combat HIV infection.

**Importance:** Studies defining effective cellular immune responses to human immunodeficiency virus (HIV) and simian immunodeficiency virus (SIV) have largely focused on a rare population that express specific MHC class I alleles and control virus replication in the absence of antiretroviral treatment. This leaves in question whether similar effective immune responses can be achieved in the larger population. The majority of HIV-infected individuals mount CD8+ T cell responses that target variable viral regions that accumulate high-frequency escape mutations. Limiting T cell responses to these variable regions and targeting invariant viral regions, similar to observations in rare ‘elite controllers’, may provide an ideal strategy for the development of effective T cell responses in individuals with diverse MHC genetics. Therefore, it is paramount to determine whether T cell responses can be redirected towards invariant viral regions in individuals without ‘protective’ MHC alleles and if these responses improve control of virus replication.

## Introduction

During HIV infection, virus-specific CD8+ T cell responses are associated with resolution of peak viremia. These responses exert substantial immune pressure that often results in rapid selection for viral escape variants, suggesting that limiting viral escape is beneficial and may prove critical in the design of immunotherapies for HIV (1–4). There is evidence that the control of viremia associated with individuals expressing specific ‘protective’ major histocompatibility complex (MHC) class I alleles may be attributed to CD8+ T cells that target specific peptide epitopes within highly invariant regions where mutations are likely to impart a significant fitness cost (5–8). However, this rare population of ‘elite controllers’ is estimated at less than 1% of the infected population, while the majority of HIV-infected individuals do not express ‘protective’ MHC alleles and more frequently mount CD8+ T cell responses that target viral regions that tolerate escape mutations easily (9, 10). Recently, multiple lines of evidence also indicate a nontraditional cytolytic role of HIV-specific CD4+ T cells that cooperate with HIV-specific CD8+ T cells to mediate suppression of virus replication and may be predictive of disease outcome (11–13). It is essential for the design of vaccines and therapeutics to determine if virus-specific CD8+ and/or CD4+ T cell responses can be mounted by individuals not expressing ‘protective’ MHC alleles, and if these responses are effective at controlling viremia. For an intervention to be truly effective, a universal approach that can contend with the extraordinary sequence diversity of HIV in people with and without ‘protective’ HLA alleles is needed.

Using nonhuman primates, we can determine if acute-phase T cell responses targeting invariant viral regions can control primary viremia. Similar to humans, some macaques express ‘protective’ MHC class I alleles associated with control of SIV replication (14–16). However, these studies have yet to define why control of virus replication in individuals expressing ‘protective’ MHC alleles is incompletely penetrant, and they do not address how to induce viral control in animals without ‘protective’ MHC alleles (17). Far less is known about the specificity of SIV-specific CD4+ T cell responses and whether they may also directly suppress virus replication. Only recently, has there been interest in developing immunogens to elicit antiviral T cells targeting conserved viral regions across individuals with diverse MHC alleles, *in vivo* (18–20). Mauritian cynomolgus macaques (MCMs) are ideal for studying pathogen-specific T cells because they have extremely restricted MHC class I and II genetics, such that nearly all of their MHC alleles can be explained by 7 common haplotypes, termed M1-M7 (21). As a result, MHC-identical animals with the potential to present identical T cell peptide epitopes can be selected for studies (21, 22).

Our group and others have reported that M3/M3 MCMs poorly control infection with pathogenic SIVmac239, making them a good example of individuals with ‘non-protective’ MHC alleles in which to characterize favorable immune responses that could be elicited in a greater proportion of the population (23, 24). Unlike pathogenic SIVmac239, replication of live-attenuated SIVmac239Δnef is controlled in nearly every infected animal, regardless of host MHC genetics. Control of SIVmac239Δnef replication in a host with ‘non-protective’ MHC alleles may be a more favorable environment in which to find the characteristics of effective immune responses that control pathogenic virus replication in the broader population. Therefore, this unique model of MHC-independent control in M3/M3 MCMs may allow the characterization of effective T cell responses in animals without ‘protective’ MHC alleles.

Previously, our group reported data suggesting that control of SIVmac239Δnef relied on immunodominant CD8+ T cell responses that select for escape mutations (25). However, at the time of our previous study, the CD8+ T cell responses restricted by MCMs expressing the M3 haplotype were incompletely known and no SIV-specific M3-restricted CD4+ T cell responses were identified. Additionally, the m3KOΔnef virus used in that study included additional mutations outside of known M3-restricted epitopes with unknown impacts on virus replication (25). We wanted to improve upon the m3KOΔnef virus by creating a virus where only known epitopes were disturbed, and mutations in other regions of the virus were avoided. Since that time, we have improved our understanding of M3-restricted CD8+ T cell epitopes and now know 10 epitopes in SIVmac239 that select for high frequency mutations (22) (25–27).

In the current study, we used this new information to create a variant of SIVmac239Δnef, termed SIVΔnef-8x, that ablated the eight M3 MHC class I-restricted epitopes that accumulate mutations during infection with SIVmac239Δnef. We hypothesized that limiting the development of CD8+ T cell responses targeting highly variable epitopes may promote the development of alternate T cell responses that target invariant regions to suppress SIVmac239Δnef replication in animals with ‘non-protective’ MHC class I alleles. We identified six immunogenic regions in SIVΔnef-8x whose immunogenicity had not previously been defined in SIV-infected M3/M3 MCMs. Four of these regions did not accumulate mutations, despite eliciting detectable responses. Interestingly, all four invariant regions were restricted by M3 MHC class II molecules and were made exclusively by animals that controlled replication of SIVΔnef-8x. These data suggest that viral control is achievable in animals with ‘non-protective’ MHC alleles even when immunodominant CD8+ T cell responses that are normally elicited during SIVmac239Δnef infection are absent. Our findings provide support for the inclusion of immunogens able to elicit CD4+ T cell responses that target invariant viral antigens in the design of an effective HIV vaccine, as this approach may be applied for widespread use across individuals with a diverse array of MHC genetics.

## Results

**Construction of SIVΔnef-8x**. We engineered a variant of SIVmac239Δnef with point mutations in eight CD8+ T cell epitopes restricted by MHC class I molecules expressed by the M3 haplotype (Figure 1a). All eight epitopes are highly immunogenic and accumulate high frequency escape mutations in response to immunodominant CD8+ T cell responses elicited during infection with either SIVmac239 or SIVmac239Δnef (16, 26, 27). For each epitope, we used sequence data from nine M3/M3 MCMs chronically infected with SIVmac239 to identify common variants in the replicating virus population (26). We performed IFNγ ELISPOT assays with the variant peptides of each of the 8 epitopes. Using PBMC collected from several SIV-infected M3/M3 MCMs, we identified variant peptides for each epitope that elicited a significantly lower IFNγ ELISPOT response *in vitro* compared to responses against the corresponding wild-type peptide (data not shown). We then used either these identified variant peptide sequences or the variant peptide sequences we included in m3KOΔnef into SIVmac239Δnef (25). We referred to the resulting virus as SIVΔnef-8x to denote its origin as an SIVmac239Δnef derivative with point mutations aimed at disrupting the immunogenicity of the eight variable M3-restricted CD8+ T cell epitopes present in SIVmac239Δnef (Figure 1a). To test if the epitope variants incorporated into SIVΔnef-8x affected viral fitness, we performed *in vitro* co-culture competition assays with a barcoded SIVmac239Δnef (BCVΔnef) containing 10 synonymous changes in *gag* that can be detected by a separate qPCR assay (25) (28). Using different ratios of barcoded virus (BCVΔnef) relative to the query virus (SIVmac239Δnef or SIVΔnef-8x), our data indicated that the eight variant epitopes we incorporated into SIVΔnef-8x did not substantially alter viral fitness *in vitro* (Figure 1b).

**Figure 1:**
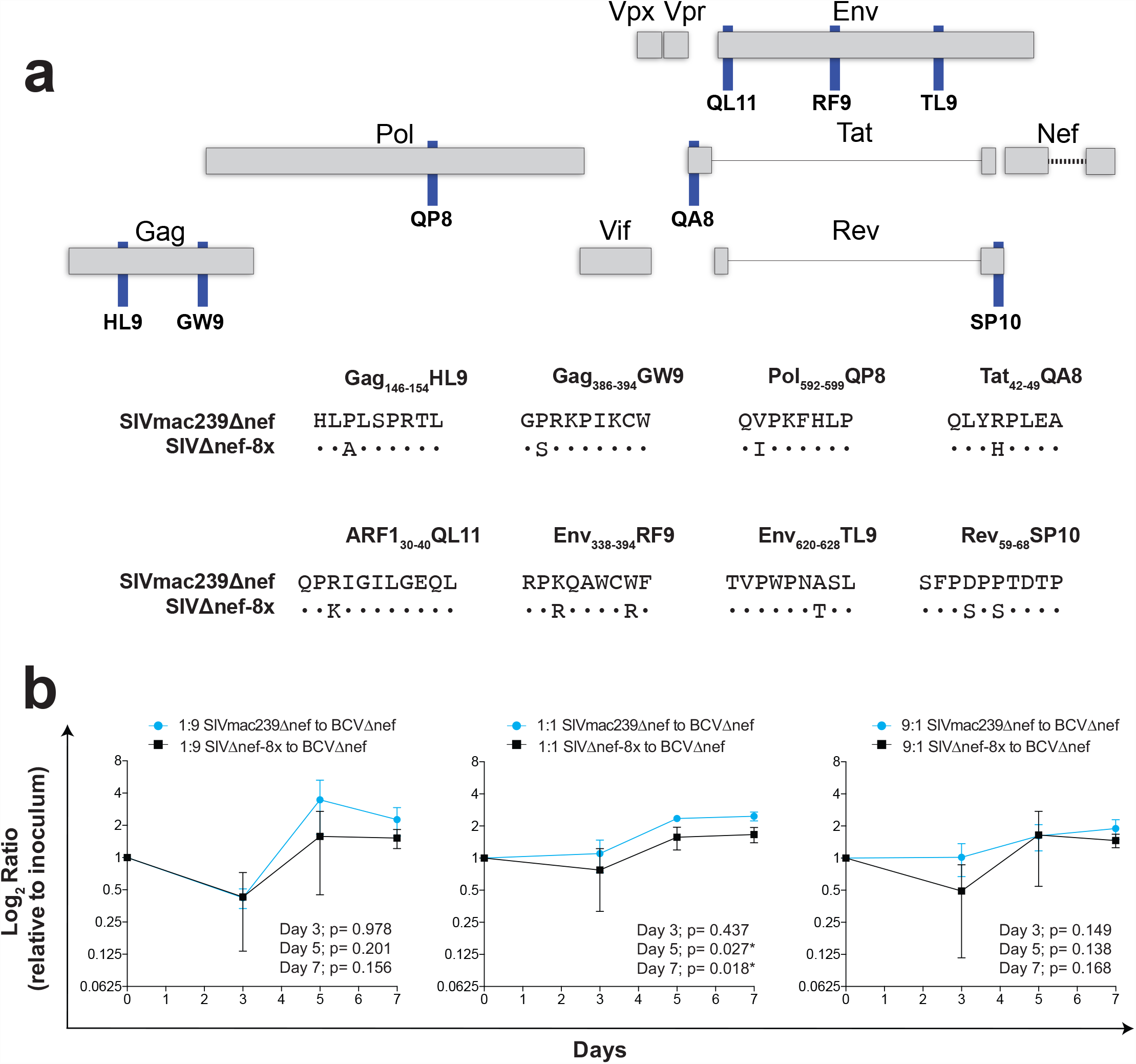
Construction of SIVΔnef-8x. (a) Diagram of live-attenuated SIV with epitope locations (top) and amino acid sequence differences (bottom) between SIVΔnef-8x and SIVmac239Δnef. Point mutations were engineered into 8 M3-restricted epitopes known to accumulate high frequency mutations during SIV infection. (b) SIVΔnef-8x and SIVmac239Δnef in vitro co-culture fitness assay. For each assay, we included a barcoded wild-type SIVmac239Δnef (BCVΔnef) as a reference at a 1:9, 1:1, or 9:1 ratio of ng p27 content relative to the query virus, either wild-type SIVmac239Δnef or SIVΔnef-8x, in the inoculum. The number of copies of query virus and barcoded virus was determined by qRT-PCR at each timepoint. A ratio of the number of query virus to barcoded virus was made and compared to the ratio present in the inoculum. All data represent mean with standard deviation of triplicate values. Unpaired t tests were performed at each time point. (*) p-value < 0.05.

**Engineered variant epitopes in SIVΔnef-8x prevent the development of responses to wild-type epitopes and are minimally immunogenic *in vivo***. We infected eight M3/M3 MCMs with SIVΔnef-8x, six M3/M3 MCMs with SIVmac239Δnef, and three M4/M6 MCMs with SIVΔnef-8x (Table 1). The T cell responses present in M4/M6 MCMs infected with SIVΔnef-8x should mirror those that would develop in M4/M6 MCMs infected with SIVmac239Δnef as these animals do not express any of the alleles of the M3 MHC haplotype (21). To determine if the mutations we incorporated into SIVΔnef-8x were sufficient to prevent the development of expected responses targeting the eight wild-type epitopes, we performed IFNγ ELISPOT assays using wild-type and variant peptides with PBMC collected throughout infection. At 3 weeks post infection, PBMC from all six M3/M3 animals infected with SIVmac239Δnef recognized up to five of the eight M3-restricted wild-type epitopes (Figure 2a, blue). In contrast, only one of eight M3/M3 animals infected with SIVΔnef-8x had positive IFNγ ELISPOT responses for any of the wild type epitopes (Figure 2a, black). None of the M4/M6 MCMs infected with SIVΔnef-8x made responses specific for the eight wild-type M3-restricted epitopes (data not shown). Similar results were observed in all animals at 8 weeks post SIVΔnef-8x infection (Figure 2b, black) and 12 weeks post SIVmac239Δnef infection (Figure 2b, blue). Therefore, M3/M3 MCMs infected with SIVΔnef-8x did not develop the previously reported CD8+ T cell responses that target variable epitopes during SIVmac239Δnef infection.

**Table 1.** Animals used in this study.

| Animal ID | Gender | MHC Haplotype | Infecting Virus |
| --- | --- | --- | --- |
| cy0684 | female | M3/M3 | SIVmac239Δnef |
| cy0686 | female | M3/M3 | SIVmac239Δnef |
| cy0687 | male | M3/M3 | SIVmac239Δnef |
| cy0689 | male | M3/M3 | SIVmac239Δnef |
| cy0749 | male | M3/recM1M3* | SIVmac239Δnef |
| cy0752 | female | M3/recM2M3* | SIVmac239Δnef |
| cy0748 | male | M4/M6 | SIVΔnef-8x |
| cy0751 | female | M4/M6 | SIVΔnef-8x |
| cy0754 | male | M4/M6 | SIVΔnef-8x |
| cy0685 | female | M3/M3 | SIVΔnef-8x |
| cy0688 | male | M3/M3 | SIVΔnef-8x |
| cy0690 | male | M3/M3 | SIVΔnef-8x |
| cy0750 | male | M3/M3 | SIVΔnef-8x |
| cy0753 | female | M3/M3 | SIVΔnef-8x |
| cy0755 | female | M3/M3 | SIVΔnef-8x |
| cy0756 | female | M3/M3 | SIVΔnef-8x |
| cy0757 | female | M3/M3 | SIVΔnef-8x |

\*Expressed the major MHC class I A and B alleles present in the M3 MHC haplotype, but also expressed minor MHC class I A alleles of the M1 (cy0749) and M2 (cy0752) MHC haplotypes. Transcriptionally abundant (major) MHC-I transcripts are responsible for restricting SIV-specific CD8<sup>+</sup> T cell responses {Budde et al., 2011, #38444} and both animals expressed the major MHC class I A allele present in the M3 haplotype (A1\*063). Accordingly, we include these functional M3/M3 animals in our group of M3/M3 MCMs infected with SIVmac239Δnef.

**Figure 2:**
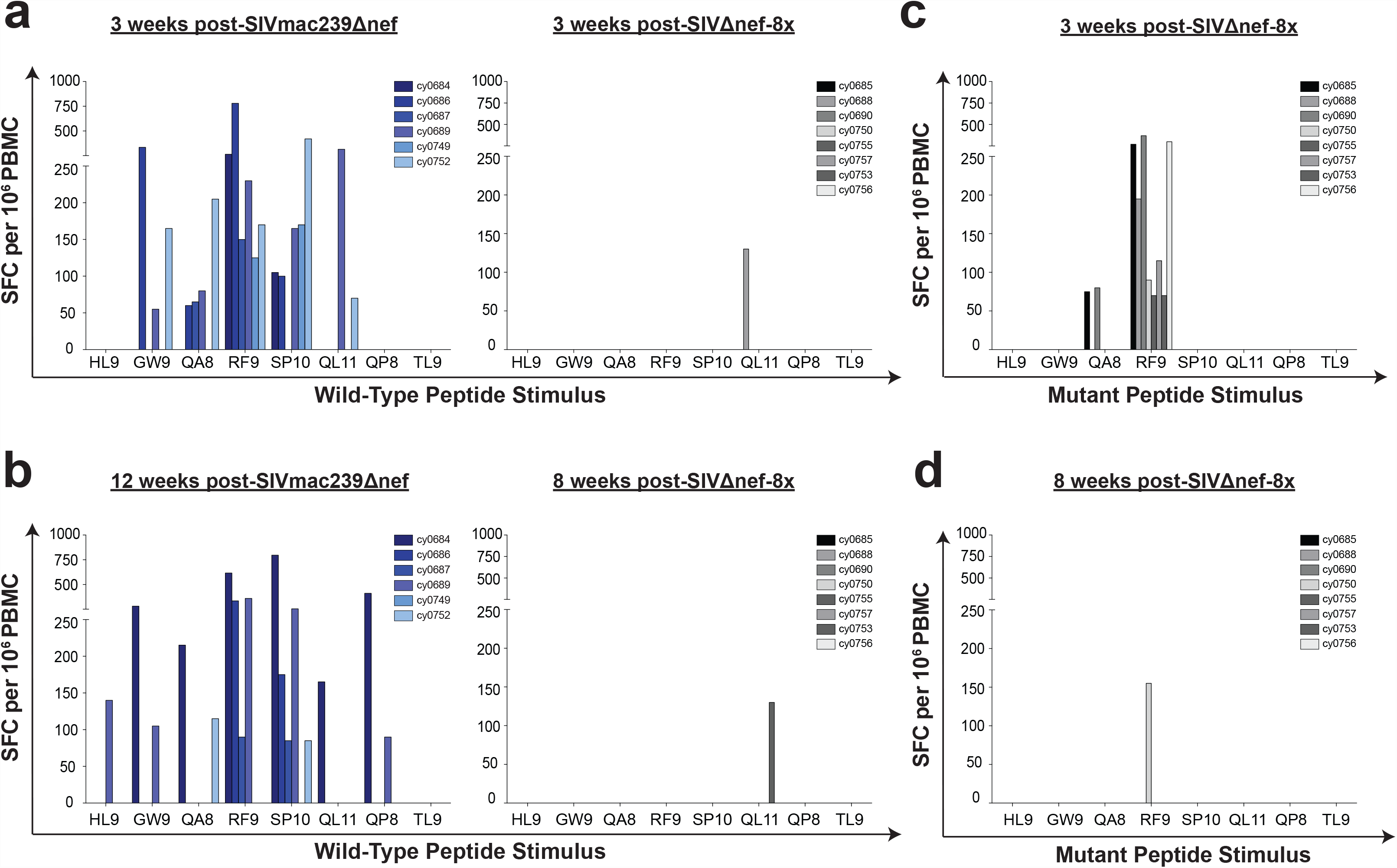
M3-restricted CD8+ T cell responses elicited during SIVmac239Δnef infection are silenced in M3/M3 MCMs infected with SIVΔnef-8x. IFNγ ELISPOT assays using peptides that match the wild-type epitope sequences were performed at week 3 (a) and week 8 or 12 (b) post infection with SIVmac239Δnef (blue) or SIVΔnef-8x (black). IFNγ ELISPOT assays using peptides that match the variant epitope sequences present in SIVΔnef-8x were performed at week 3 (c) and week 8 (d) post SIVΔnef-8x infection.

To determine if the variants incorporated in SIVΔnef-8x were immunogenic, we performed parallel IFNγ ELISPOT assays using peptides that matched the variant epitope sequences engineered into SIVΔnef-8x. At 3 weeks post SIVΔnef-8x infection, we observed positive IFNγ ELISPOT responses to only two of eight variant epitopes (Figure 2c). One variant epitope, Env_338–346_RF9 K3R/W8R, was immunogenic in all eight animals at 3 weeks post infection. Another variant epitope, Tat_42–49_QA8 R4H, elicited a response in two animals. These responses were diminished 8 weeks post infection, with only one animal detecting the variant epitope Env_338–346_RF9 K3R/W8R (Figure 2d).

**Majority of M3/M3 MCMs control SIVΔnef-8x**. Following infection, we assessed plasma viremia in M3/M3 MCMs infected with SIVΔnef-8x or SIVmac239Δnef, and in M4/M6 MCMs infected with SIVΔnef-8x (Figure 3a). The limit of detection for the viral load assay was 100 copies/mL. We did not observe peak viral load to be significantly different (p=0.158) in M3/M3 MCMs infected with SIVΔnef-8x or SIVmac239Δnef. While peak viremia of SIVΔnef-8x was slightly lower (p=0.046) in M4/M6 MCMs (n=3) compared to M3/M3 MCMs (n=8), it was not significantly different (p=0.275) from SIVmac239Δnef in M3/M3 MCMs (n=6) (Figure 3b). Taken together, these results strongly suggest that the variants incorporated into SIVΔnef-8x did not drastically alter fitness, *in vivo*. We also evaluated set point viral load, calculated as the geometric mean of viral loads between 14 and 30 weeks post infection with SIVmac239Δnef or SIVΔnef-8x (Figure 3c). Five of six M3/M3 MCMs infected with SIVmac239Δnef established a set point viral load at or near undetectable levels (median=104 vRNA copies/mL), while one animal (cy0687) had a set point viral load of 18,300 vRNA copies/mL. In contrast, virus levels by week 30 were more diverse in SIVΔnef-8x-infected M3/M3 MCMs and ranged from nearly undetectable to over 7,000 vRNA copies/mL (median=468 vRNA copies/mL). Virus replication was controlled below 1,000 vRNA copies/mL in six animals, four of which kept a set point viral load below 250 vRNA copies/mL. In contrast, two animals (cy0690 and cy0755) failed to control replication during the chronic phase of infection and had circulating virus levels that exceeded 3,500 vRNA copies/mL. Set point viral load was undetectable in two M4/M6 MCMs and fluctuated around 1,000 vRNA copies/mL in one M4/M6 MCM. Set point viral load comparisons revealed no significant differences between groups.

**Figure 3:**
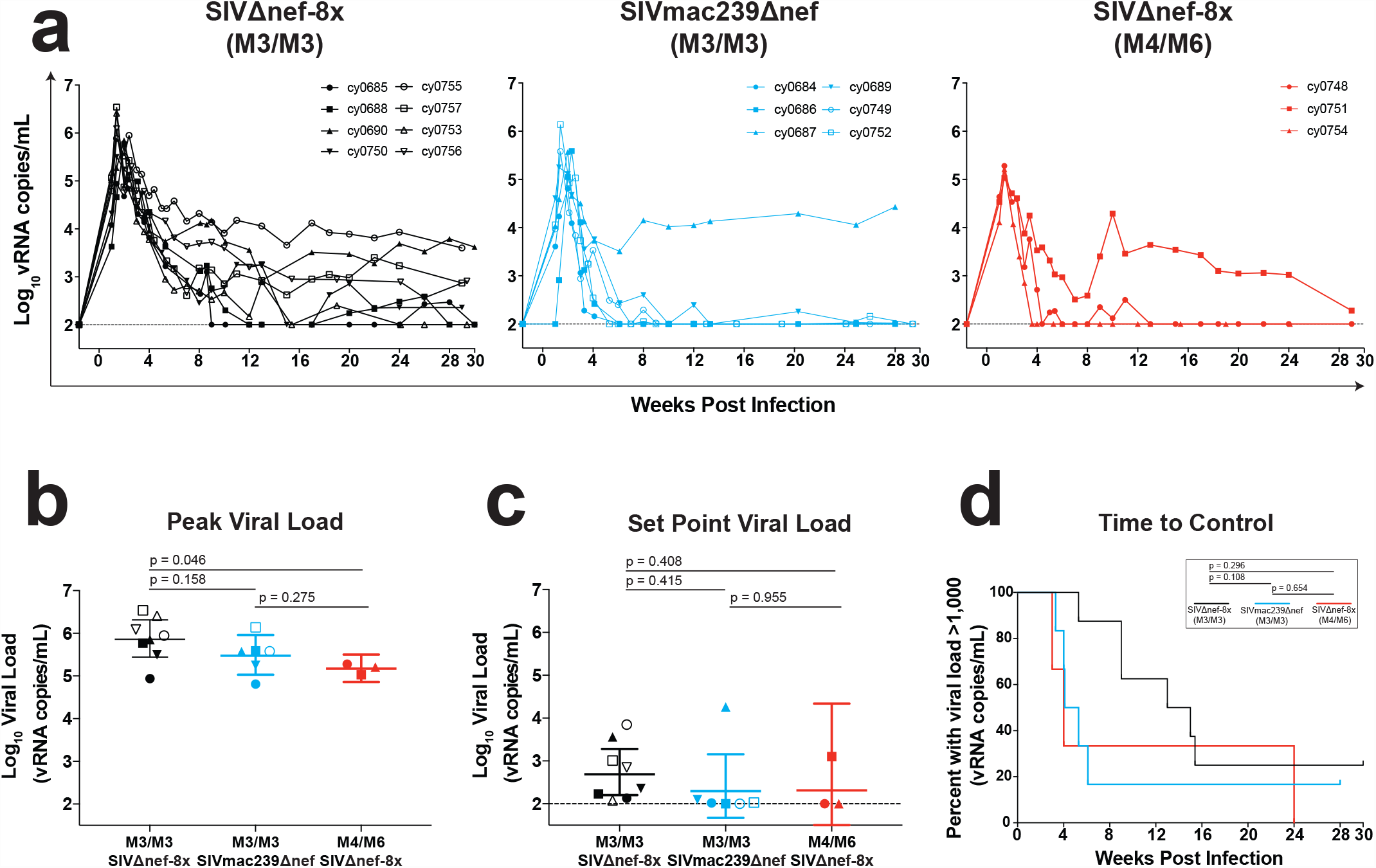
M3/M3 MCMs control SIVΔnef-8x and SIVmac239Δnef similarly *in vivo*. (a) Log10viral load trajectories were measured for 30 weeks after infection: M3/M3 MCMs infected with SIVΔnef-8x (black), M3/M3 MCMs infected with SIVmac239Δnef (blue), and M4/M6 MCMs infected with SIVΔnef-8x (red). (b) Individual peak viral load in animals infected with SIVΔnef-8x or SIVmac239Δnef displayed as geometric mean with 95% confidence intervals. (c) Comparisons of set point viral load (geometric mean of viral loads between 14 and 30 weeks post infection). (d) Initial time to control virus replication below 1,000 copies/mL. An unpaired Student’s t-test was used to calculate statistics for peak viral load and set point viral load. A log-rank test was used to calculate the statistics between groups for time to control.

Next, we compared the time to control virus replication in M3/M3 MCMs infected with SIVΔnef-8x to M3/M3 MCMs infected with SIVmac239Δnef and M4/M6 MCMs infected with SIVΔnef-8x (Figure 3d). We modeled this with Kaplan-Meier survival curves using a threshold for control similar to that of ‘elite controller’ macaques infected with SIVmac239 (<1,000 vRNA copies/mL)(14) (15). Similar to comparisons of set point viral load, we did not find significant differences in time to establish control of virus replication between the three cohorts.

**Acute-phase T cell responses elicited in M3/M3 MCMs infected with SIVΔnef-8x target several viral regions that do not accumulate mutations**. To determine if there were T cell responses present in M3/M3 MCMs infected with SIVΔnef-8x that targeted previously undefined T cell epitopes, we performed full proteome IFNγ ELISPOT assays with overlapping peptide pools to scan the entire SIVmac239 proteome. Positive peptide pools were then deconvoluted to assess responses to individual peptides. During acute (week 3) and post-peak (week 7–9) infection, we identified six immunogenic regions in the SIVΔnef-8x proteome that elicited T cell responses to peptide sequences in Gag (n=5) and Env (n=1) that were not previously characterized in SIV-infected M3/M3 MCMs. During acute infection, all six animals that controlled SIVΔnef-8x replication responded to at least one of these new regions. Four out of these six animals made responses against 3 or more of the new regions (Figure 4a, left). We observed similar results at 8 weeks post infection, where five out of the six animals controlling virus replication made one or more new responses (median=2) (Figure 4a, right). No responses were detected to any of these new regions during acute or post-peak infection in the two animals (cy0690 and cy0755) that did not control SIVΔnef-8x replication.

**Figure 4:**
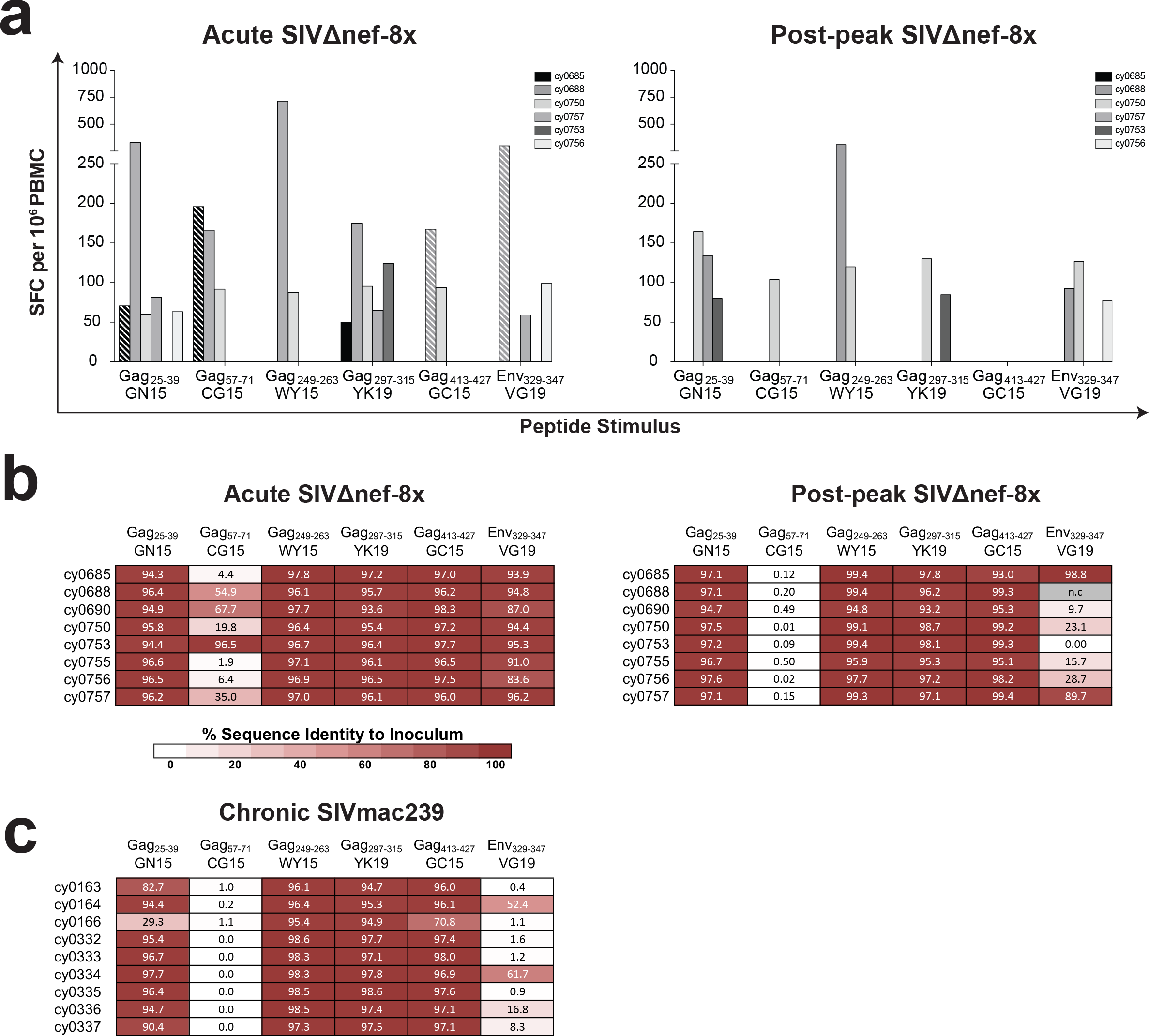
Majority of newly targeted regions do not accumulate high-frequency mutations during SIV infection. Full proteome IFNγ ELISPOT assays performed at an (a) acute timepoint (week 3) and a post-peak timepoint (week 7–9). Peptide pools were deconvoluted to assess responses to individual peptides, resulting in the identification of responses targeting six viral regions previously undefined in M3/M3 MCMs. Only positive responses are shown, with solid bars representing assays using freshly isolated PBMC and striped bars using frozen PBMC. (b) Virus populations were deep sequenced from plasma of M3/M3 MCMs during acute and post-peak infection with SIVΔnef-8x. (c) The six regions were also analyzed in M3/M3 MCMs chronically infected with SIVmac239 (>52 weeks) from a previous study by our group. Heat maps represent the percent sequence identity to inoculum. Darker colors correspond to a higher sequence identity. n.c. no sequence coverage

We deep sequenced viral populations in parallel to determine if these six newly targeted regions accumulated point mutations. During acute infection only one of the six new regions, Gag_57–71_CG15, accumulated high frequency mutations in a majority of animals (Figure 4b, left). When we examined the sequences from virus populations isolated during post-peak infection (weeks 7–9), four of the six new regions remained nearly identical to the inoculum, while Gag_57–71_CG15 and Env_329–347_VG19 had accumulated high-frequency mutations (Figure 4b, right). To determine whether these 4 apparently invariant regions can accumulate mutations during pathogenic SIV infection, we examined variant accumulation in these 4 regions in virus populations isolated and sequenced from 9 M3/M3 MCMs who had been infected with SIVmac239 for ∼52 weeks for a previous study by our group (Figure 4c) (26). In 3 of these regions, we found that 70–99% of sequences (median=97%) matched wild type SIVmac239 in all nine animals. In eight animals, more than 80% of Gag_25–39_GN15 sequences matched wild-type SIVmac239, while one animal had only 30% of sequences that matched wild-type SIVmac239. Taken together, our data suggests that it is possible to exert viral control when animals develop acute-phase T cell responses targeting regions that do not readily accumulate mutations.

**CD4+ T cells targeting regions that do not accumulate mutations are common during SIVΔnef-8x infection**. We wanted to determine whether CD4+ or CD8+ T cells were responsible for targeting the six immunogenic regions identified in the IFNγ ELISPOT assays. We grew T cell lines specific for these six peptides, and mapped MHC restriction. Of the four invariant viral regions that elicited responses, Mafa-DRA*01:02:01/DRB1*10:02 restricted both Gag_249–263_WY15 and Gag_297–315_Y19, while Mafa-DPA1*13:01/DPB1*09:02 restricted both Gag_25–39_GN15 and Gag_413–427_GC15 (Figure 5a). MHC restriction was also mapped for the two responses targeting viral regions that accumulated high-frequency mutations during SIVΔnef-8x infection. Using the optimal peptides contained within the regions, we found that Gag_57–71_CG15 was restricted by Mafa-B*011:01, while Env_329–347_VG19, was restricted by Mafa-A1*063:02 (Figure 5b). A summary of these newly targeted regions and their restricting alleles is shown in Table 2. Thus, acute phase CD4+ T cells targeting viral regions that do not accumulate mutations are common during acute infection in animals without favorable MHC genetics that ultimately control virus replication.

**Table 2.**
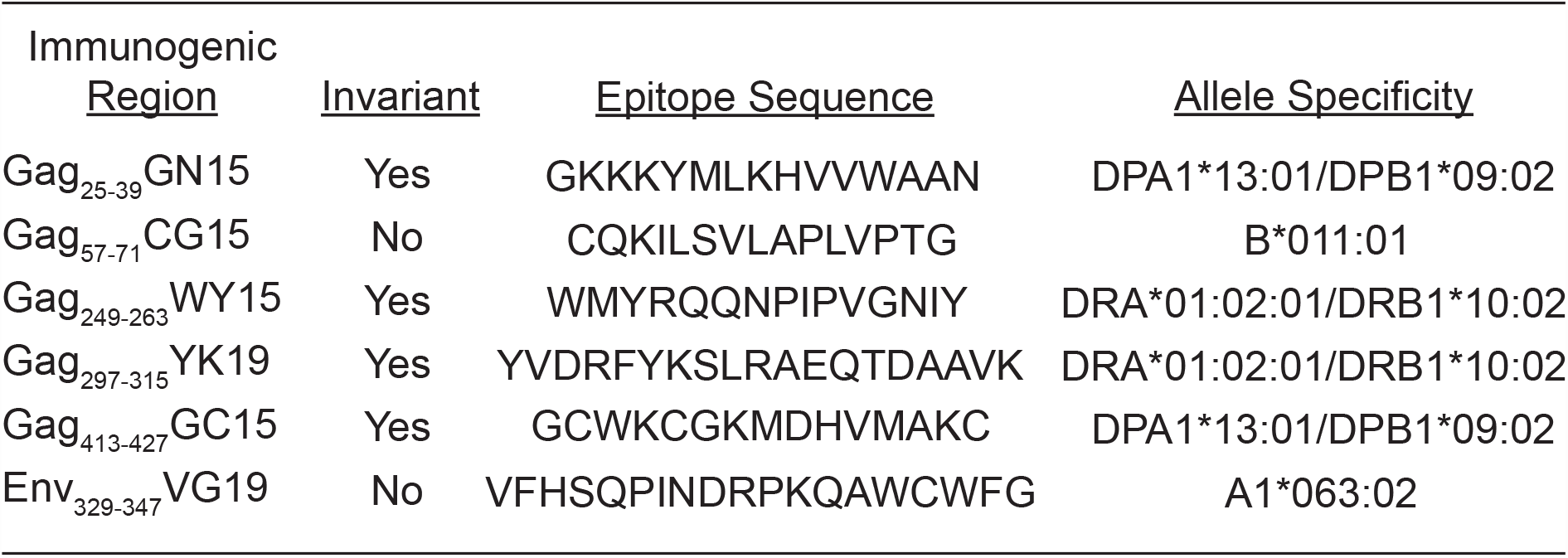
Characterization of M3-restricted SIVΔnef-8x T cell responses

**Figure 5:**
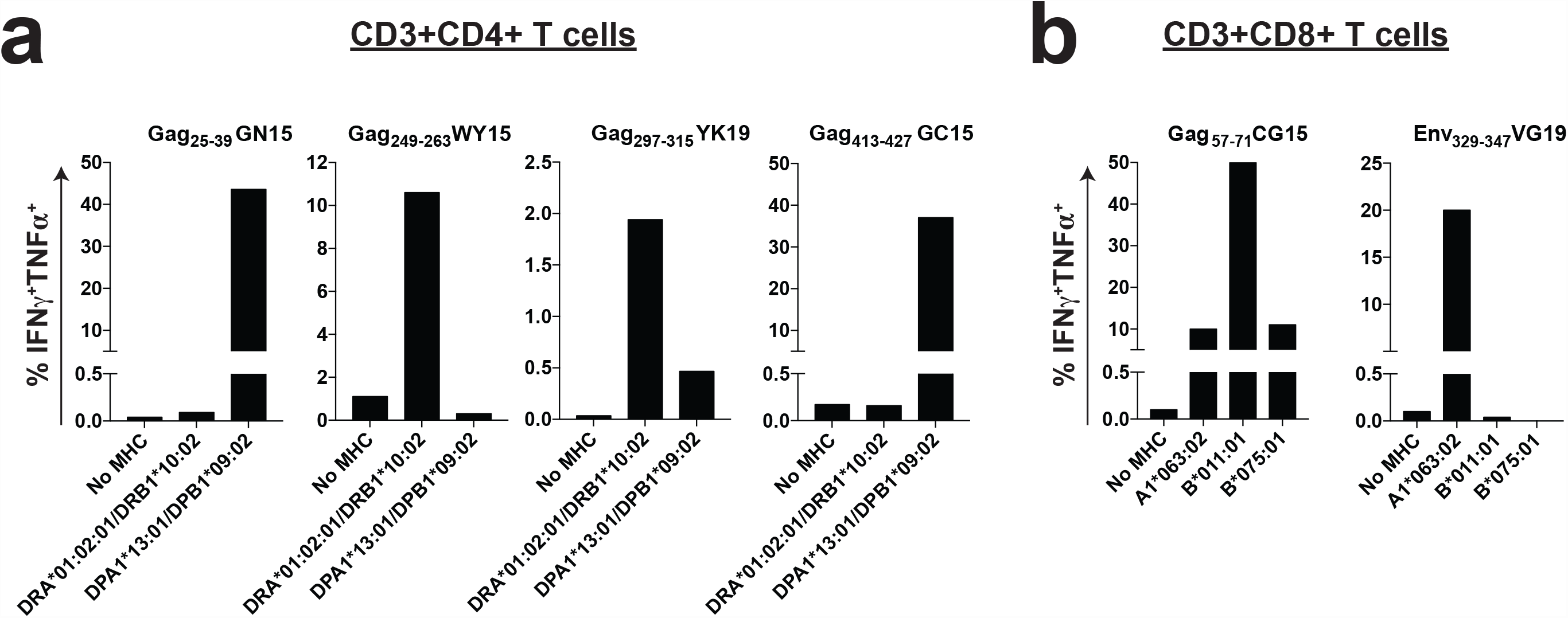
Characterization of M3-restricted SIVΔnef-8x T cell responses. ICS assays were performed to determine (a) MHC class II restriction for CD4+ T cell lines specific for the four identified invariant regions and (b) MHC class I restriction for CD8+ T cell lines specific for the two identified variable regions. Data represents percentage of cells positive for IFNγ and/or TNFα for each T cell line. MHC-matched 721.221 cells or K562 cells expressing the indicated M3 MHC class I alleles, or RM3 cells expressing the indicated M3 MHC class II alleles were used as antigen-presenting cells. Peptide-pulsed untransfected 721.221 cells, K562 cells, or RM3 cells were used as

## Discussion

Rare individuals that express ‘protective’ MHC alleles control HIV replication without antiretroviral treatment and often make CD8+ T cell responses that target highly invariant, possibly evolutionarily conserved, viral regions (7, 8, 37). In contrast, responses to invariant viral regions are frequently subdominant in HIV-infected individuals that do not express ‘protective’ MHC alleles (10). It is hypothesized that an effective vaccine will need to limit T cell responses made to highly immunogenic variable regions, in order to maximize the likelihood of developing T cell responses against invariant viral regions (38–40). Using SIVΔnef-8x, we directly tested this hypothesis in M3/M3 MCMs that do not express ‘protective’ MHC alleles and observed that control of SIVmac239Δnef can still be achieved.

Cellular responses that target Gag during acute HIV infection have been associated with low viral load and improved disease outcome (2, 8, 9). Out of the six regions that elicited T cell responses in M3/M3 MCMs infected with SIVΔnef-8x, five were located within Gag. All six animals that controlled SIVΔnef-8x replication made T cell responses during acute infection that recognized one or more of these five regions that span the majority of Gag: Gag_249–263_WY15 and Gag_297–315_YK19 are located within the p27 capsid (CA), Gag_57–71_CG15 and Gag_25–39_GN15 are located within the p15 matrix (MA), and Gag_413–427_GC15 is located within p6. Of note, the first 9 amino acids of Gag_297–315_YK19 are present in an evolutionarily conserved motif known as the major homology region (MHR) and share three of the four residues identified as highly conserved within the MHR (41). The MHR is also contained within the C-terminal subdomain of CA (CA-CTD), one of only two regions of HIV-1 Gag that is absolutely required for assembly, suggesting this region may represent an ideal target for T cell-based vaccines (42). In all six animals that controlled SIVΔnef-8x, four of the five Gag regions that were targeted did not accumulate mutations. Interestingly, the same four regions did not accumulate high-frequency mutations in nearly all nine M3/M3 MCMs chronically infected with SIVmac239 (>52 weeks) that were sequenced by our group for a previous study (Figure 4c) (26).

Besides the four invariant regions, we found two regions that accumulated mutations during acute SIVΔnef-8x infection. Gag_57–71_CG15, accumulated both a Q58R and V63A mutation by 3 weeks post infection that was present at a frequency of 7%-37% and 4%-88%, respectively. By 9 weeks post infection, less than 1% of the circulating virus matched the inoculum in Gag_57–71_CG15 for all eight M3/M3 MCMs infected with SIVΔnef-8x (Figure 4b, right). Even though the V63A mutation in Gag_57–71_CG15 was observed coincident with breakthrough viremia in an ‘elite controller’ rhesus macaque infected with SIVmac239 (43), we found the V63A mutation was present in virus populations from both controllers and non-controllers of SIVΔnef-8x (data not shown). Our results suggest that this mutation, alone, does not confer breakthrough replication, so perhaps targeting this region of Gag may still offer some immunological benefit.

We also observed high-frequency mutations in Env_329–349_VG19; a region that is located at the C-terminus of the V3 loop (44). This region also contains the variant epitope Env_338–346_RF9 K3R/W8R that we engineered into SIVΔnef-8x and was immunogenic during acute infection in all eight animals (Figure 2c). The most common mutation we observed within the newly targeted Env_329–349_VG19 was an R to W change at position 345 of Env that restores one of the two mutations in the Env_338–346_RF9 back to the original SIVmac239 sequence. It is possible that the R to W reversion was a consequence of the development of T cells targeting Env_338–346_RF9 K3R/W8R during acute infection. Alternately, it is possible that tryptophan at position 345 may be more favorable than the arginine we engineered into the virus. Given the proximity of this arginine to the C311-C344 link that serves as the base of the V3 loop, it seems possible that the polarity of the amino acid at position 345 may impact the formation of the V3 loop. The relatively conserved nature of the V3 loop, as well as its role in determining coreceptor tropism, further support this region as a site for effective T cell responses to exert their antiviral function during acute infection (44–46).

There is growing evidence that HIV-specific CD4+ T cells may play a bigger cytolytic role in control of virus replication than previously thought (11, 37, 47, 48). SIV-specific CD4+ responses have been previously detected in macaques infected with pathogenic SIV (49, 50). While observed to be typically subdominant in comparison to many CD8+ T cell responses, immune pressure imparted by CD4+ T cells has been sufficient to select for escape variants in certain cases (28, 43, 51). We found four immunogenic regions that elicited acute phase CD4+ T cells in M3/M3 animals infected with SIVΔnef-8x. All four of these were located in Gag and did not accumulate mutations by 9 weeks post infection. Notably, the same regions of Gag did not routinely accumulate mutations in M3/M3 MCMs infected with SIVmac239 for a year (Figure 4c). To our knowledge, this is the first identification of MHC class II-restricted SIV-specific T cell responses in MCMs. Although we were unable to dissect a mechanism of CD4+ T cell mediated control of virus replication during acute SIVΔnef-8x infection, cytolytic CD4+ T cells have been previously implicated in control of viral infections (11, 47, 52). This argument lies in contrast to virus-specific CD4+ T cells as preferential targets of infection, and may be explained by distinct transcriptional and functional signatures of cytolytic CD4+ T cells that mirror CD8+ T cells more than Th1 CD4+ cells (13, 53, 54). Together, we provide evidence that acute phase CD4+ T cells may improve control of SIV replication, and we provide a model in which to further explore this mechanism in future studies.

We did find that two M3/M3 MCMs infected with SIVΔnef-8x (cy0690 and cy0755) and one M3/M3 MCM infected with SIVmac239Δnef (cy0687) were unable to control virus replication, similar to that seen in the animals of the Harris et. al. study. Both cy0690 and cy0755 did not have T cell responses detectable by IFNγ ELISPOT assays. When we sequenced virus populations isolated from cy0690 and cy0687, we found an additional deletion in *nef* that has previously been shown restore the reading frame and result in increased pathogenicity (data not shown) (55). This observation prompted us to re-examine the sequences of virus populations replicating during chronic infection from the M3/M3 MCMs of the Harris et. al. study. We found similar sequence changes within the same region of *nef* in virus populations isolated from these animals (data not shown). The precise mechanism by which *nef* is restored in certain animals and whether functional advantages are conferred remains to be determined and requires further investigation.

Together, our study suggests that perhaps expanding subdominant virus-specific CD4+ T cells towards invariant viral regions during early infection may improve viral control. Even though not every animal in our study was able to mount these responses and elicit viral control, our data provides compelling evidence that CD4+ T cell responses targeting MHC class II-restricted epitopes have the potential to be effective and can be mounted by individuals without ‘protective’ MHC alleles. Future nonhuman primate studies may consider focusing on effective SIV-specific CD4+ T cell responses and evaluating their direct contribution to controlling virus replication *in vivo*.

## Materials and Methods

**Animal care and use**. Seventeen MCMs were purchased from Bioculture Ltd and were housed and cared for by the Wisconsin National Primate Research Center (WNPRC) according to protocols approved by the University of Wisconsin Graduate School Animal Care and Use Committee. Animals were chosen based on expression of particular MHC alleles as previously described (29–33). Eight M3/M3 MCMs and three M4/M6 MCMs were infected intravenously with 10ng of p27 SIVΔnef-8x. Four M3/M3 MCMs and two functional M3/M3 MCMs were infected intravenously with 10ng p27 of SIVmac239Δnef. Four of these MCMs were infected as part of a previous study (34), and two animals (cy0749 and cy0752) were functionally M3/M3 (Table 1).

**Creation of virus stocks**. We created plasmids (pUC57) containing the 5’ and 3’ viral genomes of SIVmac239Δnef by custom gene synthesis (GenScript, Piscataway, NJ) as done previously (25). For SIVΔnef-8x, ten substitutions were then incorporated into SIVmac239Δnef by site-directed mutagenesis (GenScript, Piscataway, NJ). The plasmids containing the 5’ and 3’ halves of the corresponding genomes were digested with SphI, treated with antarctic phosphatase, precipitated, and ligated together. Vero cells were transfected with the ligated products and co-cultured with CEMx174 cells for 48 hours. Infected CEMx174 cells were grown for ∼2 weeks to produce high-titer viruses and harvested daily during the last week. Plasma SIV loads were determined by qRT-PCR and the p27 content of the virus stocks was determined by enzyme-linked immunosorbent assay (ELISA; ZeptoMetrix Corp., Buffalo, NY) according to the manufacturer’s protocol. The p27 content of SIVmac239Δnef was 327 ng/mL, and the viral load was 2.19e9 copies/mL. The p27 content of SIVΔnef-8x was 245 ng/mL, and the viral load was 1.39e9 copies/mL.

***In vitro*** **co-culture competition fitness assays**. In triplicate, SIVmac239Δnef or SIVΔnef-8x were mixed with BCVΔnef at p27 content ratios of 1:1, 1:9, and 9:1. Each virus mixture was incubated with 1e6 CEMx174 cells at 37°C for 4 hours. After washing, 5e5 cells were plated and grown for 1 week, with supernatant being sampled at days 3, 5, and 7. A discriminating qPCR assay was used to quantify the copies of SIVmac239Δnef or SIVΔnef-8x virus and the BCVΔnef virus, as done previously (28) (25). Briefly, viral RNA (vRNA) was isolated from the inoculum and each supernatant and then quantified with the SuperScript III Platinum One-Step quantitative PCR kit (Invitrogen, Carlsbad, CA). In one reaction, primers and probes targeting an 84-bp region of *gag* were used to quantify SIVmac239Δnef and SIVΔnef-8x. A separate reaction with a distinct set of primers and probes was used to quantify BCVΔnef. The p27 content ratio of SIVΔnef-8x or SIVmac239Δnef to BCVΔnef in each supernatant was normalized to the ratio that was present in the inoculum, and replicative differences between viruses were assessed at each time point with unpaired Student’s *t* tests (GraphPad Prism, La Jolla, CA). All data represent mean with standard deviation of triplicate values and is plotted on a log_2_ scale.

**Plasma viral load analysis**. Plasma was isolated from whole blood by ficoll-based density centrifugation and cryopreserved at –80°C. SIV *gag* loads were determined as previously described (34). Briefly, vRNA was isolated from plasma, reverse transcribed, and amplified with the Superscript III Platinum one-step quantitative reverse transcription-PCR (qRT-PCR) system (Invitrogen). The detection limit of the assay was 100 vRNA copy equivalents per mL of plasma (copies/mL). The limit of detection value (100 copies/mL) was reported when the viral load was at or below the limit of detection.

**Peptides**. The NIH AIDS Research and Reference Reagent Program (Germantown, MD) provided 15-mer peptides overlapping by 11 amino acid positions spanning the full SIVmac239 proteome. Additional peptides used for mapping were created by custom synthesis (GenScript, Piscataway, NJ). All peptide sequences were derived from the SIVmac239 proteome.

**IFN-γ ELISPOT assays**. Gamma interferon (IFN-γ) enzyme-linked immunospot (ELISPOT) assays were performed using fresh and frozen PBMC as previously described (26, 34). Each of the eight variant epitopes were selected using frozen PBMC collected from at least five M3/M3 MCMs during acute or chronic SIV infection. Fresh peripheral blood mononuclear cells (PBMCs) were isolated from EDTA-anticoagulated blood by ficoll-based density centrifugation. A precoated monkey IFN-γ ELISPOTplus plate (Mabtech, Mariemont, OH) was blocked and individual peptides were added to each well at a final concentration of 1µM. Multiple peptide pools containing 15-mer peptides that span the full SIVmac239 proteome, each overlapping by 11 amino acids, were used to assess new responses that emerged during infection. Peptide-pools totaled 10µM (1µM each peptide) and were added to cells at a final pool concentration of 1µM. Each peptide or peptide-pool was tested in duplicate, as was concanavalin A (10µM) that was used as a positive control. Four to ten wells did not receive any peptides and served as a negative control for calculating background reactivity. Assays were performed according to manufacturer’s protocol and wells were imaged with an AID ELISPOT reader. Positive responses were determined using a one-tailed *t* test and an alpha level of 0.05, where the null hypothesis was that the background level would be greater than or equal to the treatment level (35). Positive responses were considered valid only if each duplicate well had a value of at least 50 SFCs per 10^6^ PBMCs. If statistically positive, reported values represent the average of the test wells minus the average of all negative control wells.

**Generation of M3/M3 BLCLs**. MHC-matched B-lymphoblastoid cell lines (BLCLs) were generated as previously described (Budde et al., 2011, #86642). Briefly, PBMCs were isolated by density-based centrifugation from whole blood containing EDTA. B cells were then immortalized with medium from an S549 cell line containing herpesvirus papio. Cells were maintained in R-10 medium and sequenced to verify the presence of appropriate MHC alleles.

**Generation of peptide-specific T cell lines**. CD8+ T cell lines were generated by incubating 5e6 freshly isolated PBMCs with 2µM peptide in complete medium (RPMI 1640 medium supplemented with 15% fetal calf serum, 1% antibiotic/antimycotic, and 1% L-glutamine) with 100 IU of interleukin-2 (NIH AIDS Research and Reference Reagent Program). The cell lines were restimulated weekly with peptide-pulsed irradiated (9,000 rads) BLCLs, as previously described (27). CD4+ T cell lines were generated by depleting freshly isolated PBMC of CD8+ cells using NHP-specific anti-CD8 microbeads via magnetic separation according to the manufacturer’s protocol (Miltenyi Biotech, San Diego, CA) Then 5e6 CD8+-depleted cells were incubated with 2uM peptide. CD4+ T cell lines were maintained similarly to CD8+ T cell lines, with media additionally containing a final concentration of 50 ng/mL interleukin-7 (BioLegend, San Diego, CA).

**Intracellular cytokine staining assay**. To determine T-cell line peptide specificity and the restricting MHC class I or class II molecule, we measured intracellular expression of interferon gamma (IFNγ) and tumor necrosis factor alpha (TNFα) as previously described (27, 36). Briefly, 2e5 peptide-pulsed BLCLs or MHC class I or class II transferents were incubated for 1.5 hrs with peptide at 37°C, washed twice with RPMI medium containing 10% FBS (R10), and then combined with 2e5 cells from corresponding CD4+ or CD8+ T cell lines for an additional 4 hrs at 37°C in the presence of brefeldin A (Sigma-Aldrich, St. Louis, MO). Cells were incubated with LIVE/DEAD fixable near-IR dead cell stain for 15 min before being surface stained with CD3-AF700 (clone: SP34–2; BD Biosciences), CD4-APC (clone: M-T466; Miltenyi Biotec), and CD8-Pacific Blue (clone: RPA-T8; BD Biosciences) for 30 min in the dark at room temperature. Cells were fixed with 2% paraformaldehyde (PFA) for at least 20 min, and then permeabilized with 0.1% saponin and stained with IFNγ (clone: 4S.B3; BD Biosciences) and TNFα-PerCP/Cy5.5 (clone: MAb11; BD Biosciences) for 30 min in the dark at room temperature. Flow cytometry was then performed on an LSR II instrument (BD Biosciences) using 2% PFA-fixed cells, and data were analyzed using FlowJo, Version 9.9.6 (Treestar, Ashland, OR).

**Deep Sequencing of SIV**. Replicating virus populations were subjected to genome-wide deep sequencing, as previously described (26). Briefly, viral RNA was isolated from plasma with the MinElute virus spin kit (Qiagen) and amplification of cDNA was generated using the Superscript III one-step reverse transcription PCR (RT-PCR) system with high-fidelity Platinum *Taq* (Invitrogen). This resulted in four overlapping amplicons spanning the entire SIV coding sequence that were then purified by the MinElute gel extraction kit (Qiagen) and quantified using the Quant-IT double-stranded DNA (dsDNA) HS assay kit (Invitrogen). Pooled amplicons (1ng) were used to generate uniquely tagged libraries using the Nextera XT kit (Illumina). In select cases, cDNA was generated using SuperScript IV First Strand Synthesis System (Invitrogen) and PCR amplified with Q5 high-fidelity DNA polymerase (NEB) in duplicate to generate 37 overlapping amplicons spanning the entire SIV coding sequence. Amplified products were quantified using the Quant-IT double-stranded DNA (dsDNA) HS assay kit (Invitrogen) and diluted to 3ng/uL for library preparation with TruSeq Nano HT (Illumina). Tagged libraries were then quantified using the Quant-IT dsDNA HS assay kit and fragment size distribution was assessed using a high sensitivity Agilent bioanalyzer chip. Libraries were then pooled and sequenced on an Illumina MiSeq.

All sequences were analyzed with Geneious (Biomatters, Ltd.). Paired reads were initially quality trimmed using BBDuk (decontamination using kmers) plugin, a part of the BBTools package by Brian Bushnell, and then mapped with high sensitivity to the SIVmac239 sequence; gaps up to 500-bp were allowed when mapping in order to identify any addition deletions or insertions (GenBank accession number M33262). Variant nucleotides were called at a threshold of 1%. The variants detected in the analyzed virus populations were then compared to the mutant sites originally incorporated into SIVΔnef-8x.

## Statistics

Student’s *t* test was used evaluate the significance of differences between peak viral load and set point viral load (geometric mean of viral loads from 15 to 30 weeks post infection).

All viral load measurements were log_10_ transformed. Kaplan-Meier survival analyses were used to model the time to control virus replication in M3/M3 MCMs infected with SIVΔnef-8x (n=8) or SIVmac239Δnef (n=6), and M4/M6 MCMs infected with SIVΔnef-8x (n=3); log-rank (Mantel-Cox) test was then used to determine significance of differences. All statistical analyses were performed using GraphPad Prism (GraphPad Prism, La Jolla, CA).

## Acknowledgements

Research reported in this publication was supported in part by the Office of The Director, National Institutes of Health under Award Number P51OD011106 to the Wisconsin National Primate Research Center, University of Wisconsin-Madison. Research reported in this publication was supported in part by National Institutes of Health Award Number R01AI108415, as well as by the National Institute of General Medical Sciences of the National Institutes of Health under Award Number T32GM081061. The content is solely the responsibility of the authors and does not necessarily represent the official views of the National Institutes of Health The content is solely the responsibility of the authors and does not necessarily represent the official views of the National Institutes of Health Special thanks to Jason Weinfurter, Matthew Reynolds, and Adam Ericsen for helpful discussion.

